# Amplitude-modulated kilohertz stimulation targeting beta-band activity disrupts motor learning

**DOI:** 10.64898/2026.07.11.737109

**Authors:** Philipp Reber, Christina M. Merrick, Guy Avraham, Ida Killebrew, Katheryn Thayer-Pham, Haajar Ahmad-Ali, Angel V. Peterchev, Karunesh Ganguly, Cidnee Luu, Daniel Sheltraw, Ludovica Labruna, Richard B. Ivry

## Abstract

Sensorimotor learning is associated with the modulation of neural rhythms in the primary motor cortex (M1). Beta-band activity (“beta”; 15–30 Hz) is suppressed during learning, while delta oscillations (1–4 Hz) become increasingly correlated with movement kinematics as skill improves. These observations have been complemented by experimental manipulations designed to perturb oscillatory activity with externally applied electric fields (E-fields). In non-human primates, invasive beta stimulation has been shown to disrupt motor learning whereas delta stimulation enhanced motor recovery in a stroke model. Causal evidence in humans remains limited, partly because established non-invasive methods cannot achieve continuous, narrowband E-fields at sufficient amplitude in the brain. To address this gap, we employed kilohertz transcranial magnetic perturbation (kTMP), a non-invasive magnetic induction technique that delivers continuous narrowband kilohertz E-fields which can be amplitude-modulated (AM) to target cortical rhythms. We applied AM-kTMP to test the functional relevance of beta and delta activity in human motor learning. In a double-blind mixed design, 40 participants performed a force-control task while receiving AM-kTMP at E-field amplitudes of 8 V/m in M1. We targeted either beta or delta, each paired with a sham condition. AM-kTMP influenced motor performance in a frequency-dependent manner: Beta-kTMP suppressed performance gains relative to both delta-kTMP and sham, whereas delta-kTMP showed no effect. These results suggest that increased beta activity in human M1 can interfere with motor learning. More broadly, kTMP offers a novel approach to probe frequency-specific cortical dynamics.

## Introduction

Learning new motor skills, whether riding a bike or playing the saxophone, requires repeated practice. The acquisition and refinement of requisite movements rely on spatiotemporal changes in brain activity across distributed cortico-subcortical circuits^1^. Within these circuits, the primary motor cortex (M1) plays a central role, exhibiting practice-dependent reorganization^2^. Changes in oscillatory dynamics provide one physiological signature of learning^3^, evident in multiple frequency bands including beta-band activity^4,5^ (“beta”; 15–30 Hz) and low-frequency oscillations^6–8^ (delta 1–4 Hz; theta 4–7 Hz). While these oscillatory dynamics are thought to help coordinate signals across regions and promote spike-timing dependent plasticity^9,10^, their precise role in driving learning remains incompletely understood.

Current insights into these dynamics stem from observations of simple or well-rehearsed complex movements. Beta-band activity typically desynchronizes across M1 preceding and during voluntary movement^11^, manifest as a transient suppression of beta power^12,13^. Following movement, beta reemerges^14,15^. This post-movement rebound is modulated by experience: it is attenuated following movement errors^5^ and elevated following well-learned or automatized movements^3,16^. Excessive beta synchronization across the motor network is associated with impaired movement initiation in Parkinson’s disease^17^ and reduced dexterity after stroke^18^.

Together, these observations support the view that synchronized (high-power) beta represents an anti-kinetic or stabilizing signal, potentially constraining plastic changes during learning^16,19^.

By comparison, the presence of low-frequency oscillations in M1 has been associated with accurate and fast motor performance. Delta oscillations encode muscle activation patterns critical for precise movement control and align with the timing of sub-movements^20–23^. Delta activity has been linked to motor learning as it emerges with task proficiency^24^, an effect hypothesized to reflect the coordination of task-relevant neural activity across brain regions^21,25^. Consistent with these findings, delta oscillations are greatly reduced in power after stroke in humans, as well as in animal models of stroke, and their reappearance is predictive of the recovery of motor function^20,21,23^. Collectively, these findings point to low-frequency oscillations as a key organizing rhythm that supports the fine temporal structure of movement that is refined during learning^25^.

To determine whether frequency-specific neural rhythms causally shape motor learning, studies have targeted motor cortex with oscillating electric fields (E-fields) and evaluated the impact on skill acquisition. In invasive studies with non-human primates, epidural alternating current stimulation (eACS) targeting beta notably disrupted performance gains on a visuomotor learning task^19^. Conversely, eACS targeting delta has been shown to enhance motor performance in reach-to-grasp tasks specifically during recovery from M1 injury, effects that were attenuated as recovery plateaued^21,25^.

Non-invasive brain stimulation (NIBS) studies in human participants present a more variable picture. Transcranial alternating current stimulation (tACS) targeting beta activity in M1 has produced inconsistent effects on motor learning, with reports of behavioral impairment^26^, facilitation^27,28^, and no detectable change^29–33^. Similarly, tACS targeting delta activity has yielded mixed results^31,34,35^. This variability may reflect at least in part the limited E-field amplitude reaching the cortex (*≤* 0.5 V/m)^36–38^, constrained by the tolerability of peripheral nerve stimulation. Repetitive transcranial magnetic stimulation (TMS), by contrast, delivers pulsed high-amplitude E-fields that can promote^39^ or disrupt^40^ plasticity depending on pulse rate. Yet, the pulsed nature of repetitive TMS induces a broadband E-field which may limit its ability to selectively interrogate frequency-specific brain dynamics. Together, such considerations highlight an opportunity to explore new NIBS approaches that can deliver continuous, narrow-band E-fields at amplitudes suitable for testing frequency-specific perturbations of neural activity^41^.

Here, we used a novel non-invasive magnetic induction technique, kilohertz transcranial magnetic perturbation (kTMP)^42,43^, during an isometric force-control task to test how frequency-specific targeting of M1 activity influences motor learning. kTMP generates a continuous narrow-band kilohertz (kHz) E-field that can be amplitude-modulated (AM) to target specific physiological rhythms. Prior work shows that AM kHz fields can modulate neural activity^43–45^ and may alter behavior^46,47^. We previously observed that kTMP targeting M1 with an E-field amplitude of 2 V/m reliably increased corticospinal excitability in three different cohorts, without causing tactile sensations^43^. In the present study, we employed AM-kTMP in a double-blind design to target either beta or delta activity in M1 with cortical E-field amplitudes of 8 V/m. Our core prediction was that beta-kTMP would disrupt motor learning. Moreover, we predicted that this effect would be frequency-specific, with delta-kTMP instead sparing learning and potentially enhancing performance.

## Results

Participants (*N* = 40) performed an isometric force-control task (Figure 1A) while receiving active kTMP in one session and sham stimulation in a second session, with the order counterbalanced. Half of the sample (*n* = 20) received beta-kTMP with an AM frequency of 21.2 ± 3.0 Hz (*M* ± *SD*) determined from individual EEG recordings. The other 20 participants received delta-kTMP with the AM frequency set to 3 Hz (Figure 1C). The M1 stimulation target was functionally localized for each participant using single-pulse TMS (see <u>Methods</u>). A double-blind mixed design allowed us to assess kTMP effects relative to sham within subjects, and to evaluate the frequency-specific nature of these effects by comparing the beta-kTMP group with the delta-kTMP group. Task performance was quantified using the mean absolute error between produced and target forces, reaction time (RT), and the rate of force development (RFD), defined as the average slope from force onset to peak force. For each participant, session, mapping, and target, these metrics were normalized against the corresponding performance in the first experimental task block.

**Figure 1.**
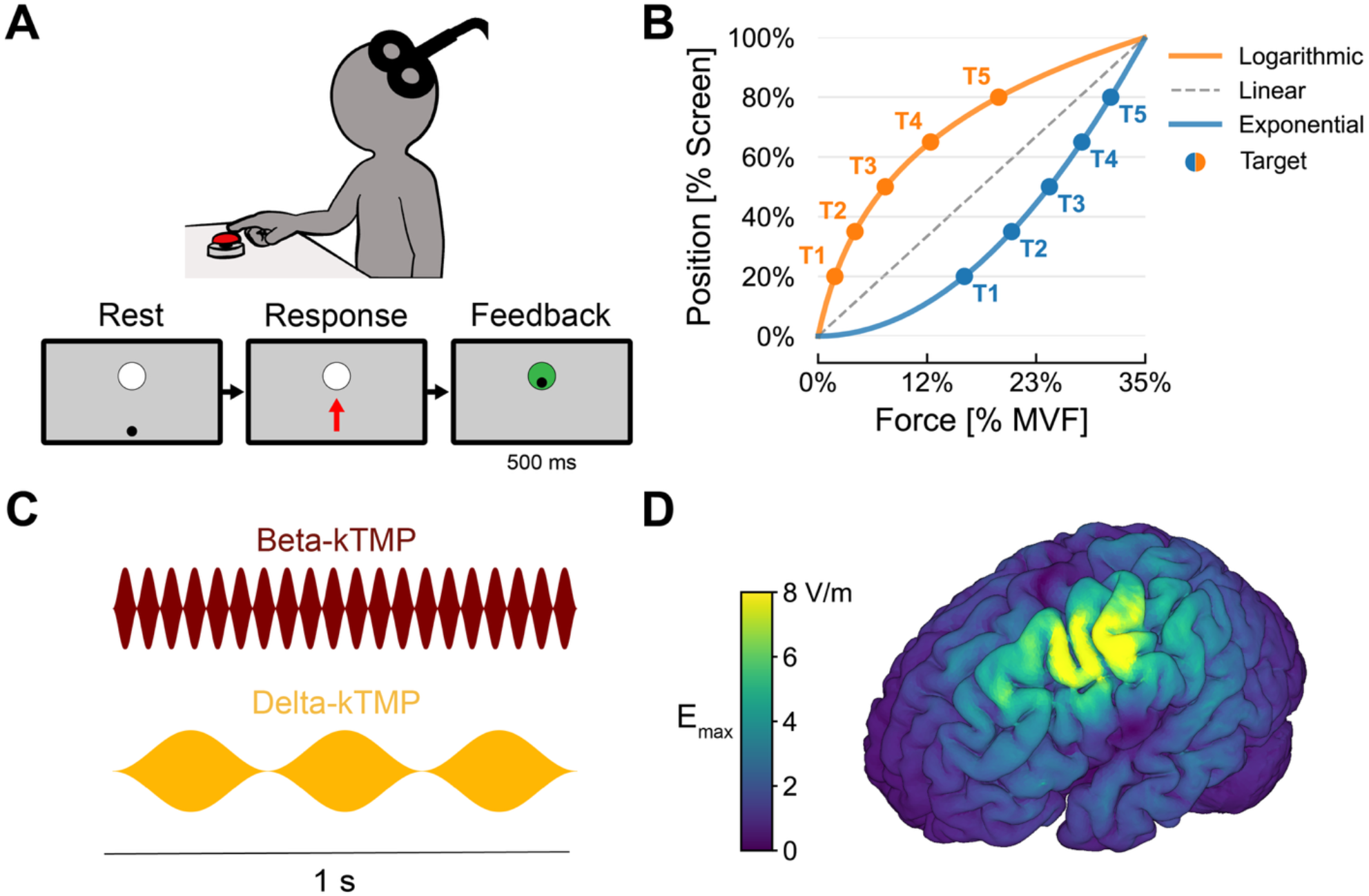
Experimental overview. **(A)** In the force control task, participants produced isometric pulses on a force sensor with their index finger, attempting to match the target force cued by a white circle that appeared at one of five vertical positions. A cursor (black dot) was visible at rest and, after each response, indicated the peak force of that pulse, with the target briefly turning green on hits and red on misses. Targets were shown in shuffled sequences of five, with each sequence including one trial of each target. Every sequence was followed by 2 s of additional performance feedback (see <u>Behavioral task</u>). **(B)** Participants completed two sessions spaced 1–2 days apart. Each session included a warm-up block using a linear force-cursor mapping (30 trials per target) followed by five testing blocks using one of two non-linear mappings (30 trials per target per block, with 30 s rest between blocks). **(C)** kTMP was continuously applied over M1 for 20 min during testing blocks and the breaks between them. kTMP parameters were set to induce a cortical E-field amplitude *E*^max^ = 8 V/m (active) in one session, and 0 V/m (sham) in the other. For 20 participants, the kTMP waveform was amplitude-modulated (AM) to match the participants’ estimated beta-band peak frequency (beta-kTMP); for the other 20 participants, the AM frequency was set to 3 Hz (fixed for all participants; delta-kTMP). **(D)** The simulated spatial distribution and magnitude of the cortical E-field induced by active kTMP targeting M1 in the left hemisphere.

### Task performance

Behavior improved over time as participants performed the task (Figure 2). This was evident in a time-dependent decrease in error (i.e. accuracy gains; linear mixed-effects model [LME], effect of Block: *β* = –0.042 [–0.058, –0.026], SE = 0.008, *t*(234) = –5.13, *p* = 6.07 × 10^−7^; see <u>Statistical analysis</u>). This gain in accuracy was accompanied by a progressive decrease in RT (*β* = –0.025 [–0.036, –0.013], SE = 0.006, *t*(235) = –4.20, *p* = 3.86 × 10^−5^; Figure S1B, top). The concurrent improvements in both accuracy and RT suggest that participants gained skill rather than simply shifting along a speed-accuracy function over time^48^.

**Figure 2.**
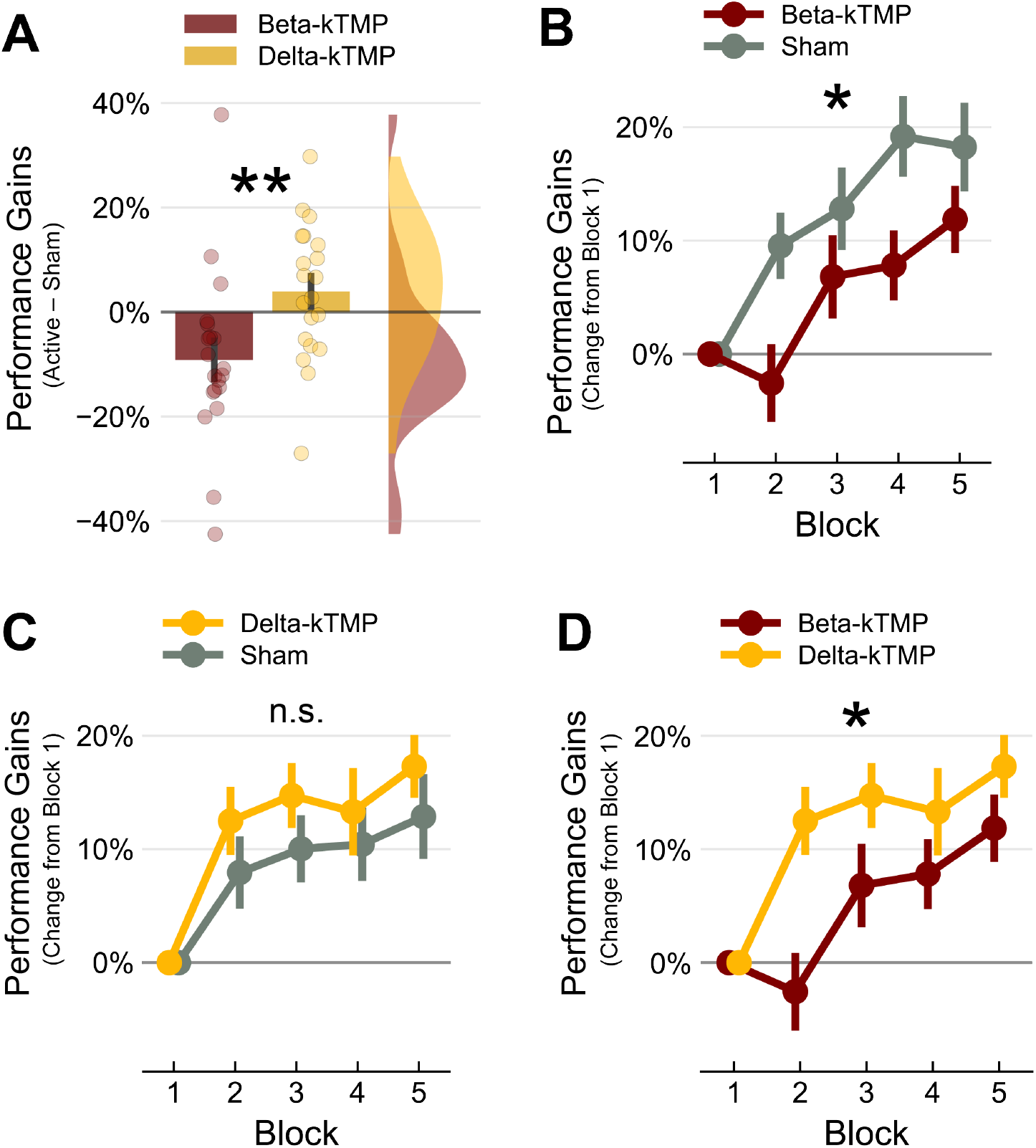
AM-kTMP produces a frequency-specific effect on motor performance gains. Performance gains are expressed as percent change in error with respect to the first block, with values sign-flipped so that positive values indicate improvement. All error bars are standard errors. **(A)** The Condition × Frequency interaction: the effect of stimulation (Active *™* Sham) attenuated performance gains in the beta-kTMP group compared to the delta-kTMP group. **(B)** Within the beta-kTMP group, performance gains under active stimulation were reduced relative to sham. **(C)** Within the delta-kTMP group, active stimulation and sham were comparable. **(D)** Under active stimulation, performance gains were lower in the beta-kTMP group than in the delta-kTMP group (data from active conditions replotted from B and C).

Per design, the distinct shapes of the two non-linear force-cursor mappings (Figure 1B) required opposite force adjustments relative to the warm-up trials. Under the logarithmic mapping, participants had to reduce the force produced for each target; under the exponential mapping, they had to increase the produced force. These changes were accompanied by corresponding decreases and increases in the RFD, respectively (LME, effect of Mapping: *β* = – 0.193 [–0.253, –0.132], SE = 0.031, *t*(36) = –6.13, *p* = 4.62 × 10^−7^; Figure S1A, left). The specific mapping also influenced overall task performance. We observed larger gains in accuracy under the logarithmic mapping than under the exponential mapping (*β* = –0.049 [–0.092, –0.006], SE = 0.022, *t*(36) = –2.21, *p* = 0.034; Figure S1A, right), while changes in RT remained comparable across mappings (*β* = –0.007 [–0.051, 0.036], SE = 0.023, *t*(36) = –0.33, *p* = 0.742).

### Personalized frequency-tuned kTMP targeting beta-band activity suppressed motor performance

Our central question was whether stimulation modulated learning and whether these effects were frequency-specific. We found that the effect of stimulation (active vs. sham) on accuracy gains depended on the targeted frequency (LME, Condition × Frequency: *β* = 0.131 [0.045, 0.216], SE = 0.045, *t*(36) = 2.90, *p* = 0.006; Figure 2A). To unpack this interaction, we first compared accuracy gains between active and sham stimulation conditions within each frequency group (beta-kTMP and delta-kTMP). Participants showed smaller gains in accuracy during beta-kTMP than during sham (*β* = 0.092 [0.027, 0.156], SE = 0.032, *t*(36) = 2.88, *p*_Holm_ = 0.027; Figure 2B). In contrast, targeting the delta band had no significant effect on behavior, as accuracy gains were similar during delta-kTMP and sham (*β* = –0.039 [–0.104, 0.026], SE = 0.032, *t*(36) = –1.22, *p*_Holm_ = 0.452; Figure 2C). We also compared the beta-kTMP and delta-kTMP groups within each stimulation condition. During active stimulation, the beta-kTMP group showed smaller gains in accuracy than the delta-kTMP group (*β* = 0.085 [0.017, 0.152], SE = 0.033, *t*(37) = 2.54, *p*_Holm_ = 0.046; Figure 2D). Importantly, accuracy gains were comparable between the subgroups during their respective sham sessions (*β* = –0.046 [–0.122, 0.030], SE = 0.037, *t*(37) = –1.23, *p*_Holm_ = 0.452).

The frequency-dependent effect of stimulation was specific to accuracy. For both RT and RFD, the effect of stimulation did not differ between the beta-kTMP and delta-kTMP groups (LME, Condition × Frequency, RT: *β* = –0.061 [–0.148, 0.026], SE = 0.046, *t*(36) = –1.34, *p* = 0.187; RFD: *β* = 0.003 [–0.120, 0.126], SE = 0.064, *t*(36) = 0.04, *p* = 0.966). Active stimulation also produced no overall change in either measure compared to sham (main effect of Condition, RT: *β* = 0.039 [–0.023, 0.100], SE = 0.032, *t*(36) = 1.20, *p* = 0.237; RFD: *β* = 0.069 [–0.018, 0.155], SE = 0.046, *t*(36) = 1.51, *p* = 0.141; Figure S1B), and the two frequency groups showed comparable changes overall (main effect of Frequency, RT: *β* = –0.021 [–0.078, 0.036], SE = 0.030, *t*(36) = –0.71, *p* = 0.480; RFD: *β* = –0.022 [–0.106, 0.062], SE = 0.043, *t*(37) = –0.50, *p* = 0.620).

### kTMP enabled effective blinding

Participants rated the intensity of scalp sensations associated with kTMP using an 11-point scale (0 = no sensation, 10 = very intense sensation) immediately after the last testing block in each session. Consistent with our informal observations, most participants reported no sensations at all, with non-zero ratings reported by only 8 of the 40 participants (20%) after active kTMP (*E*_max_ = 8 V/m) and 10 of the 40 participants (25%) after sham stimulation. The median rating was 0 for both active and sham kTMP, with no difference between the two conditions (Wilcoxon sign-rank test: *W* = 28.00, *p* = 0.647; Figure 3A). We also found no difference in ratings of pain (*W* = 1.50, *p* = 1.000), discomfort (*W* = 11.50, *p* = 0.498), annoyance (*W* = 30.00, *p* = 0.776), or muscle twitches (*W* = 39.50, *p* = 0.406) between active and sham kTMP. To assess the effectiveness of our blinding procedure without revealing the existence of a sham condition (see <u>Experimental protocol</u>), participants were asked at the end of the second session to indicate whether that session involved active or sham stimulation. Eighteen of 40 participants (45%) correctly identified their stimulation condition in the second session, a rate not exceeding chance (two-sided exact binomial test, *p* = 0.636, 95% CI [29.3%, 61.5%]).

**Figure 3.**
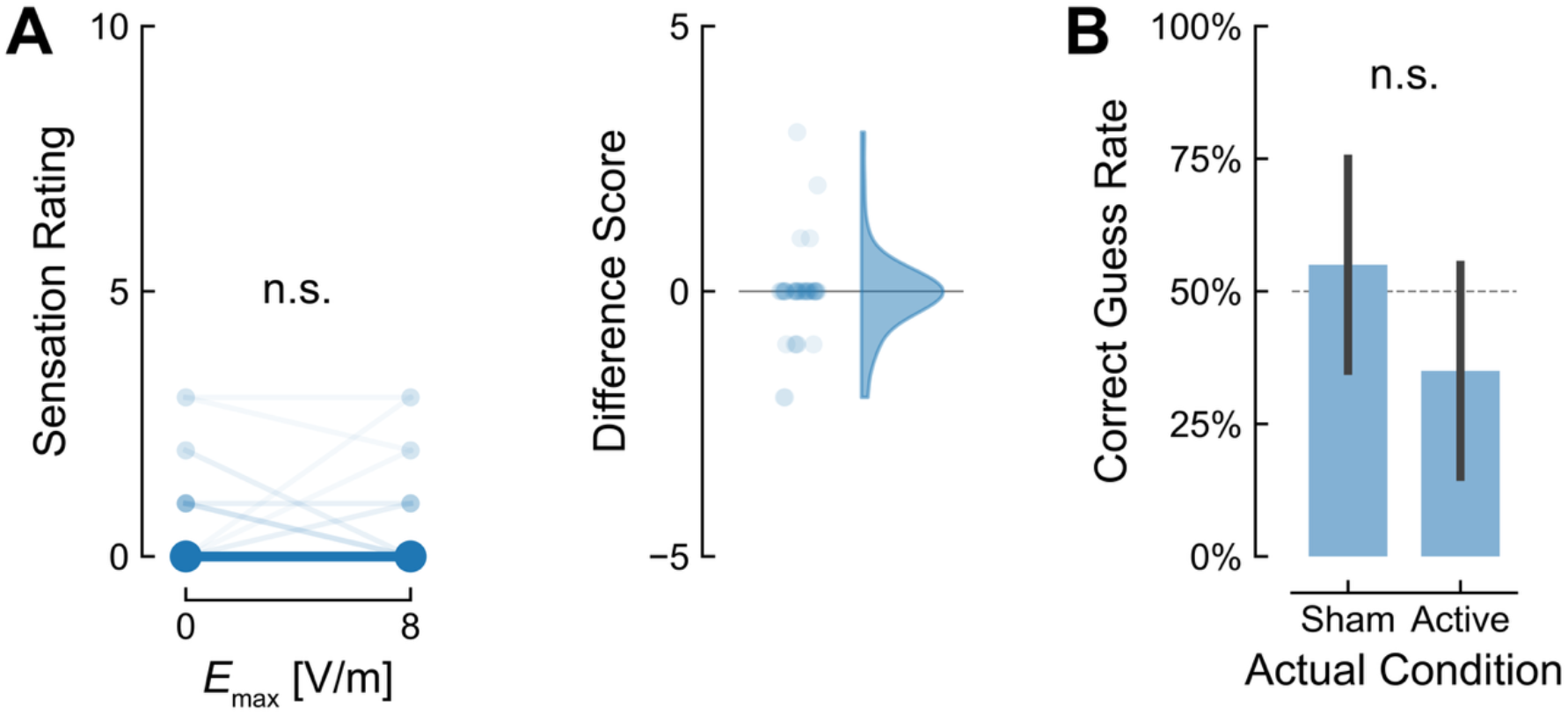
kTMP is indistinguishable from sham stimulation. **(A)** *Left:* Ratings of intensity of scalp sensation were similar following active (*E*^max^ = 8 V/m) and sham (0 V/m) AM-kTMP stimulation over M1. Large circles represent group medians. *Right:* Distribution of difference scores (Active – Sham) calculated for each individual. Saturation level of dots and lines indicate the number of observations per score. **(B)** Proportion of participants who correctly indicated the stimulation condition. Participants did not perform above chance level. Error bars indicate 95% CI (not visible in A).

## Discussion

Our results indicate that non-invasive, frequency-tuned perturbations of M1 can selectively interfere with the acquisition of a new motor skill. To induce the perturbation, we used kTMP, a novel magnetic induction technique in which a kHz carrier waveform is amplitude-modulated (AM) to target physiologically relevant frequencies. Targeting motor beta with an AM kHz field during practice of an isometric force-control task attenuated learning. This impairment was specific to the AM frequency given that delta stimulation left learning intact. The effect of beta kTMP was reflected in reduced gains in motor accuracy (error) while our other two performance metrics, reaction time (RT) and rate of force development (RFD) were not impacted by either beta or delta stimulation. These findings support the hypothesis that cortical beta-band dynamics can act as a regulator of motor plasticity in the human brain^16,19^.

Synchronized beta in M1 is thought to reflect a physiological state that favors stability over plasticity^3,5,16^. In our study, beta-kTMP applied during skill acquisition may have biased the motor network toward such a synchronized state. This bias conceivably impeded the transient beta desynchronization proposed to be linked to error processing^5^ and/or the updating of a forward model^49^. This view aligns with findings in non-human primates, where beta stimulation suppressed the reactivation of task-related neural ensembles^19^, a process that may support learning^50^.

The effect of beta-kTMP on learning only manifested as reduced gains in accuracy. Changes in RT and RFD remained similar across stimulation conditions. This observation can be interpreted through the lens of recent evidence distinguishing between global and local beta dynamics. In this framework^18^, global beta bursts reflect synchronization across cortical and subcortical regions and the strength of this activity is linked to movement inhibition and slowing, typical of Parkinson’s disease^51^ and stroke^18^. In contrast, desynchronization of local cortical beta bursts is hypothesized to support the computational flexibility required for producing complex movements^18^. The selective effect of beta-kTMP over M1 on accuracy suggests the perturbation was sufficient to disrupt local beta desynchronization associated with refining a novel movement pattern, while having a negligible effect on global beta activity that is associated with the descending drive required for movement execution. We recognize that mechanistic inferences about the behavioral dissociations must be viewed with considerable caution. Additional studies are required to assess whether the dissociation of accuracy and speed measures following cortical stimulation with kTMP replicates and generalizes across different experimental conditions and manipulations.

Our observations diverge from prior work reporting slowed movements under beta stimulation with low-amplitude tACS^52,53^. This discrepancy may stem from the distinct biophysical properties of electrical contact methods versus magnetic induction techniques. tACS delivers a sinusoidal current at the target frequency, whereas AM-kTMP delivers a 3.5 kHz sinusoidal carrier amplitude-modulated at the target frequency. Consequently, the neural elements affected by these waveforms are likely different^43^. Moreover, tACS E-field distributions are heavily constrained by the heterogeneous conductivities of scalp and skull, while the magnetic induction utilized by kTMP is largely independent of these tissue properties. This independence allowed us to explore higher cortical E-field amplitudes without eliciting sensory effects. In contrast, tACS intensities are capped near tolerability limits due to co-stimulation of the retina^54^ or peripheral nerves^55^, effects that can be addressed by titrating the current below perception thresholds or by including an appropriate control montage^56^. kTMP limits the risk of unblinding even at the high field strengths used here, a methodological consideration we have empirically verified in the current study. Recognizing these mechanistic differences is important for the comparative interpretation of stimulation effects across different neurotechnologies, and they may explain why our results differ from those of previous work with beta-tACS.

We included the delta perturbation condition given prior invasive work suggesting that delta stimulation with epidural alternating current stimulation targeting the motor network can improve motor performance^21^. However, our results showed no statistically significant differences between delta-kTMP and sham stimulation on any of the performance measures. Research involving a non-human primate stroke model suggests that performance gains from delta stimulation may be state-dependent: the benefits of delta stimulation were pronounced during early recovery but diminished as motor function was regained^21^. Furthermore, the timing of stimulation modulated these effects. Delta-driven enhancement depended on the relationship between the phase of the stimulation waveform and the temporal structure of movement^21^. The fixed, open-loop protocol used in the present study may lack the temporal precision required to reinforce phase-dependent kinematic coding. Future work should evaluate whether adaptive AM-kTMP aligning the phase of the waveform envelope with endogenous neural signals or the timing of movement can enhance performance of individuals with a healthy motor system.

To account for neuroanatomical variation, the stimulation target was localized individually for each participant based on physiological responses using a standard single-pulse TMS procedure. We chose to optimize the coil position using motor-evoked potentials from the first dorsal interosseous (FDI) even though the primary agonists for the button press are the forearm flexors (flexor digitorum superficialis and profundus). We opted for this strategy because the FDI is more reliably localized by single-pulse TMS than the forearm flexors. Given the spatial overlap of intrinsic and extrinsic hand muscle representations in M1^57^, we reasoned this would deliver a largely overlapping E-field distribution to task-relevant cortical territory. Employing the same induction coil for both localization and kTMP intervention allowed us to match the cortical E-field distribution induced during task performance to the individual functional cortical target and maintain spatial consistency across experimental sessions. We expect that this methodological choice improved the reliability of delivering the intended E-field to the motor target.

### Limitations

Several limitations of our findings warrant consideration. The behavioral task was designed to minimize interference between sessions by employing qualitatively different stimulus-response mappings to allow within-subject contrasts (active vs. sham stimulation). While this design reduced carry-over effects, it resulted in little overlap between the target forces for the two conditions and correspondingly, higher response variability for the exponential mapping due to the greater force requirements^58,59^. In addition to counterbalancing the assignment of mapping and stimulation condition, we normalized performance for each mapping, participant, and target. This approach imposes a rigorous criterion for intervention efficacy, as any performance changes evident in the initial testing block are effectively removed from the analysis.

Regarding our neural metrics, time constraints prevented the assessment of individual beta peak frequencies during the motor task itself. Our evaluation relied instead on a resting-state estimate, which may only provide a coarse approximation of the oscillatory dynamics active during motor behavior. Future work should compare endogenous frequency tuning at rest and during task performance to facilitate the development of optimized protocols for inducing neural and behavioral changes with AM-kTMP or other NIBS methods.

Finally, while participant blinding was empirically verified, experimenter blinding was only informally assessed. Although standard protocols were followed to maintain objectivity, a formal assessment for experimenter bias is important, particularly in complex stimulation setups where the experimenter must actively manage research equipment.

### Conclusion

The current study provides the first evidence that non-invasive, frequency-tuned magnetic perturbation of M1 can impact the acquisition of motor skills. Beyond the first demonstration that this novel form of NIBS can impact behavior, the disruptive effect of beta-kTMP adds further support for the hypothesis that beta-band activity serves as a regulator of motor plasticity. Moreover, the dissociation between beta-kTMP and delta-kTMP provides initial evidence that AM-kTMP can produce frequency-specific effects with high blinding efficacy.

Future work with kTMP will be essential to assess the robustness of these findings and to elucidate the underlying neural mechanisms impacted by neuromodulation with AM kHz fields.

## Methods

### Participants

Forty healthy young adults (24 female; 36 right-handed) aged 22.0 ± 3.2 years (*M* ± *SD*) completed the study. Participants were recruited from the University of California, Berkeley student community. Interested individuals were screened for contraindications to magnetic induction methods (kTMP and TMS), electromyography (EMG), and electroencephalography (EEG) prior to experimental procedures to ensure participant safety. We also used this screening to exclude participants who reported a history of neurological or psychiatric conditions, recent drug use, or current serious medical condition. Of those who met our inclusion criteria, five were replaced due to technical errors during data collection, one who failed to return for the second session, and one for dozing off despite our stimulating task. All procedures were approved by the Institutional Review Board of the University of California, Berkeley. Participants provided written informed consent and were financially compensated for their participation.

### Behavioral task

The participant was seated at a table in front of a 24-inch computer screen. Isometric force pulses were produced by applying pressure with the index finger of the dominant hand to a button-shaped force transducer. The target force for each response was cued by the vertical position of a white circle (2.7 cm diameter), with weak forces near the bottom of the screen and stronger forces near the top of the screen (Figures 1A, B). The participant was instructed to match the target force as quickly and accurately as possible by making a brief, isometric response. At rest, a cursor (a black circle, 0.4 cm diameter) was visible at the bottom of the screen which disappeared when the response was initiated. The cursor reappeared for 500 ms after the isometric pulse, with its vertical position indicating the peak force of the response.

During this interval, the target turned green to indicate a hit or red to indicate a miss.

Five target positions were spaced vertically along the central axis, at 20%, 35%, 50%, 65%, and 80% of the screen height. Cursor position was determined by one of three force-cursor mappings (Figure 1B). Under the linear mapping, the five targets required forces of 7%, 12.25%, 17.5%, 22.75%, and 28% maximum voluntary force (MVF), respectively. The logarithmic mapping corresponded to forces of 1.8%, 4.0%, 7.2%, 12.0%, and 19.3% MVF, while the exponential mapping required 15.7%, 20.7%, 24.7%, 28.2%, and 31.3% MVF. Independent of mapping, rest was defined as <0.015% MVF and the most upward position on the screen as 35% MVF.

Targets were presented in pseudo-random sequences, with each target appearing exactly once per sequence, constrained so that neighboring targets did not appear in immediate succession more than once. Following each sequence, the participant received performance feedback for 2 s indicating whether their overall response time and error for that sequence had increased or decreased relative to the median of the previous five sequences (or all available sequences at the start of the testing block). Depending on the change in speed and accuracy, the feedback read “Great! You improved”, “You got faster but less precise”, “You got more precise but slower”, or “Oh no! Less precise and slower”.

The task was structured in blocks. Each block required 150 responses, with each target presented 30 times. In each session, the participant first completed a warm-up block under the linear mapping to familiarize themselves with task, device, and feedback. Training lasted 4.5 ± 0.7 min (*M* ± *SD*). After a 1-min break, the participant completed five testing blocks. In one session, testing blocks were performed using the logarithmic mapping, while in the second session, the exponential mapping was set. These non-linear mappings increased task difficulty and, by changing the mapping between sessions, challenged the participant to acquire a new mapping during each session. A 30-second resting period was enforced between testing blocks. The total duration of the five testing blocks including rest periods was 21.9 ± 1.6 min.

### kHz transcranial magnetic perturbation (kTMP)

kTMP is a non-invasive magnetic induction technique that can be used to induce continuous kHz E-fields in tissue near the induction coil. Here, we employed the Magnetic Tides Gen3 kTMP system (Magnetic Tides Inc., Berkeley, CA, USA) consisting of a high-current switch-mode amplifier driving a MagVenture Cool B-70 figure-of-eight coil (MagVenture A/S, Farum, Denmark). By using magnetic induction, kTMP overcomes the tolerability limitations of high-current transcranial electrical stimulation, enabling larger cortical E-field amplitudes while keeping scalp exposure comparatively low. Power constraints^43^ require that the system operate in the kHz range in order to generate these E-fields at the cortical surface. In contrast to the brief, pulsed output waveform of conventional TMS, the kTMP waveform can be continuous.

The waveform can be delivered either as an unmodulated kHz field or amplitude-modulated to target physiologically relevant frequencies.

In the present study, the carrier frequency of the kTMP waveform was set to 3.5 kHz, as previous work demonstrated a neural response at this frequency^43^. To target beta or delta activity in M1, the carrier signal was amplitude-modulated with either a beta or delta envelope, yielding the following E-field waveform:

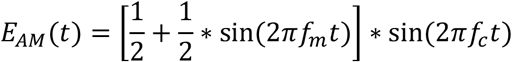

where *f*_*m*_ is the modulation (envelope) frequency, *f*_*c*_ the carrier frequency and *E*_*AM*_ the resulting time-varying amplitude-modulated waveform (Figure 1B).

kTMP parameters were set to deliver a peak current of *I* = 246 A at *f*_*c*_ = 3.5 kHz to the coil, resulting in a temporal derivative of the current *dI*/*dt* = 5.41 A/µs. *dI*/*dt* was estimated to achieve an E-field amplitude of 8.0 V/m at a depth of 14 mm perpendicular to the coil surface, approximating a location on the surface of M1 beneath the scalp^60^. Individual anatomy was not considered. The spatial distribution of the induced E-field is determined by the coil geometry and position.

#### Localization

We used a standard single-pulse TMS protocol to identify the optimal coil position relative to the head to engage our cortical target, the hand region in M1 contralateral to the response hand. The TMS pulses were delivered using a MagVenture MagPro R30 to drive the Cool-B70 coil, the same coil used for kTMP. The coil was positioned tangentially to the scalp with the handle pointing posterior-laterally at approximately 45° to the midline, inducing a posterior–anterior current flow in the underlying cortex. The stimulation target (‘hotspot’) was defined as the coil position that reliably elicited motor evoked potentials (MEPs) in the first dorsal interosseous (FDI) muscle. To this end, the coil was iteratively re-positioned over the scalp while pulse intensity was slowly increased to identify the position that produced the largest and most consistent MEPs in the FDI at minimal pulse amplitude. The hotspot was recorded using the BrainSight neuronavigation system (Rogue Research, Montreal, Canada) to ensure consistent targeting throughout the application of kTMP.

#### Electromyography (EMG)

MEPs were recorded with surface EMG (Bagnoli-8 EMG System, Delsys Inc., Natick, MA, USA). Delsys Bipolar DE-2.1 surface electrodes were placed over the belly of the first dorsal interosseous muscle (FDI) and the abductor digiti minimi (ADM) of the response hand. The ADM signal was monitored during the localization procedure to ensure that MEPs were mainly produced in the FDI, not in the ADM. A reference electrode was placed on a nearby bony landmark (lateral epicondyle or ulnar styloid process). Electrode sites were prepared by gently abrading the skin with NuPrep abrasive gel (Weaver and Company, Aurora, CO, USA) and then cleaning with isopropyl alcohol to reduce impedance. Signals were amplified, hardware band-pass filtered between 20–450 Hz and digitized at 2 kHz.

### Electric field modeling

E-field simulations were performed using the finite element method (FEM) implemented in SimNIBS (v4.5)^61^ on the default “Ernie” head model provided with the software. The MagVenture Cool-B70 coil model was positioned tangentially over C3 according to the 10-20 system layout^62^, oriented toward the contralateral orbit. Coil placement was adjusted to minimize the distance between the coil center and the cortical target while preventing intersection with the head model. The rate of change of the coil current was set to *dI*/*dt* = 5.41 × 10^6^ A/s, matching the kTMP device setting. Isotropic electrical conductivities were assigned to all tissue compartments using the default values implemented in SimNIBS. To minimize the influence of numerical artifacts at tissue boundaries, the peak E-field magnitude was defined as the 99.9^th^ percentile of the field distribution. As simulations relied on a generic rather than individualized head model, cortical E-field amplitudes are expected to vary between participants.

### Electroencephalography (EEG)

For the participants in the beta-kTMP group, EEG was used to identify the AM-kTMP envelope frequency, operationalized at the individual level as the frequency exhibiting peak power in the beta range (15–30 Hz) during 5 min of wakeful rest. EEG was recorded with a BioSemi ActiveTwo system (BioSemi B.V., Amsterdam, Netherlands) with Ag/AgCl active electrodes at a sampling rate of 8192 Hz. The amplifier was DC-coupled (input range ±262 mV; 24-bit resolution; 31 nV/bit), and Common Mode Sense and Driven Right Leg electrodes were used instead of a conventional ground. Electrodes were placed over motor cortex contralateral to the dominant hand (C1, C3, FC1, CP1; or C2, C4, FC2, CP2). The scalp was lightly abraded with NuPrep gel before electrode placement. DC offsets were kept within ±20 mV for all channels.

The EEG data were processed offline using the MNE-Python toolbox^63^. The raw time series were band-pass filtered between 1–35 Hz using a zero-phase finite impulse response filter and referenced to the common average. Ocular artifacts were removed using independent component analysis (ICA) by excluding components correlated with electrooculographic (EOG) activity detected at channel FC1/FC2. Processed timeseries were segmented into consecutive 4-second epochs, excluding those exceeding 50 µV from further analysis. Power spectral density was estimated using Welch’s method with a frequency resolution of 0.25 Hz. Spectral parameterization was performed using the FOOOF toolbox^64^ to separate periodic components from the aperiodic background. The dominant rhythm within the beta band was identified as the center frequency of the strongest oscillatory peak in the model fit. In cases where peak prominence was limited, the frequency was determined by the highest local power density within the range of interest.

### Experimental protocol

Participants were sequentially allocated to one of two groups: an initial cohort (*n* = 20) was assigned to the beta-kTMP group, followed by a second cohort (*n* = 20) assigned to delta-kTMP. Experimental conditions remained identical across both testing periods. Each participant completed two sessions spaced 1–2 days apart, one with active stimulation and one with sham stimulation. The order of the two sessions was counterbalanced, and within each group, the assignment of the two force-cursor mappings was counterbalanced.

At the beginning of the first session, individual experimental parameters were determined. For participants in the beta-kTMP group, we first completed the resting state EEG protocol to estimate the participant’s beta-band peak frequency. We tuned the AM frequency of the kTMP output waveform to the individual peak frequency aiming to facilitate perturbation effects through network resonance^65,66^. EEG was not recorded for the delta-kTMP group because endogenous delta oscillations can be difficult to distinguish from aperiodic background activity in this frequency range^67^. Rather than individualize the AM frequency for this group, we set the frequency parameter to 3 Hz based on prior research^21^.

For all participants, we used single-pulse TMS to define the cortical target for kTMP stimulation and recorded the associated coil position (“hotspot”). Following this, the participant performed a series of isometric button presses with the index finger of their dominant hand under the instructions to exert maximal force. Maximum voluntary force (MVF) was operationalized as the mean of the median forces obtained during three 3-second exertions^68^ and was set on an individual basis.

After completing the preliminary procedures, the coil was positioned on the hotspot using the neuronavigation system and remained in place for the duration of the behavioral task. During the warm-up block, sham kTMP (*E*_max_ = 0 V/m) was applied to familiarize the participant with the experimental setup. During the subsequent testing blocks (including the rest periods between these blocks), active kTMP (8 V/m) or sham kTMP was applied continuously for 20 mins, starting with task onset.

The kTMP system generates an auditory tone during active stimulation. The following steps were taken to avoid unblinding due to the auditory difference between kTMP conditions. An audio file reproducing the frequency spectrum emitted during active kTMP was played during both active and sham kTMP conditions. To further mask auditory percepts, the experimenter wore over-ear noise-attenuating earmuffs and the participant wore noise-attenuating foam-tip in-ear monitors (ER-2 Tubephone, Etymotic Research, Elk Grove Village, IL). Finally, white noise was delivered via the participant’s in-ear monitors at a level calibrated to mask all sound. A coded input system was used such that the experimenter was unaware of the kTMP condition during testing blocks.

At the end of the session, the participant completed a brief survey regarding their subjective experience of kTMP stimulation. They were asked to rate overall scalp sensations associated with kTMP stimulation on an 11-point numeric rating scale (0 = no sensation, 10 = very intense sensation). They were instructed to disregard any sensations related to the static contact of the coil with the scalp. The survey also included ratings on three questions regarding the subjective experience of pain, annoyance, and muscle twitches, using the same 11-point scale.

The second session consisted of a brief TMS procedure to verify the hotspot recorded during the first session, followed by the warm-up and testing blocks of the behavioral task, and post-stimulation survey. At the end of the second session, the participant was informed that the kTMP stimulation had been active in one session and sham in the other session. The participant was asked to indicate whether they had received active or sham stimulation in the current (second) session. We did not ask this at the end of the first session to avoid revealing the existence of a sham condition.

### Behavioral analysis

Behavioral data were preprocessed and summarized using custom-written Python code. We excluded trials in which the participant made more than one press to reach the target before a full release or when technical errors occurred (1.1% of all trials). In addition, 0.77% of data points were removed by applying a centered rolling median absolute deviation (MAD) filter (window = 10 trials). Outliers were defined as values exceeding 3 robust standard deviations (estimated as 1.4826 × MAD) from the local median^69^. To prevent the exclusion of minor deviations during periods of low variability, points were only removed if they also exceeded the 85th percentile of absolute deviations across all testing blocks. The first two trials of each block were exempted from exclusion to account for initial adaptation, thus preserving gradual changes across trials. After cleaning, the data were aggregated per participant, session, block, and target. Data distributions were tested for normality and summarized with means when normally distributed and medians otherwise.

To quantify performance gains, behavioral measures were expressed as the percent change in performance relative to the first testing block within each participant, session, and target. This normalization controlled for individual differences in baseline performance and allowed for the evaluation of relative improvements across targets and force-cursor mappings. Normalized performance changes were averaged across targets to obtain block-level performance metrics per participant. For each testing block and condition, data points exceeding three standard deviations from the mean were removed (0.17%). The remaining data were used to fit the linear mixed-effects models described below.

### Statistical analysis

Statistical analyses were performed in R (version 4.4.2) using the *lme4*^70^ package. Behavioral outcomes were modeled with linear mixed-effects models (LMEs) including stimulation Condition (Active vs. Sham), Frequency (Beta vs. Delta), Session (Session 1 vs. Session 2), Mapping (Logarithmic vs. Exponential), and Block as fixed effects and Participant as random effect. Nested models were fitted under maximum likelihood and evaluated using likelihood-ratio χ^*2*^ tests to determine the optimal model structure^70^. Three-way interaction terms involving Condition × Frequency were assessed, but only Condition × Frequency × Block improved model fit and was retained. The final model comprised all main effects, the retained interaction terms, a random intercept for Participant, and a random by-participant slope for Condition. Model diagnostics were verified visually using the R package *performance*^71^.

Estimated marginal means and pairwise contrasts were computed with *emmeans*^72^. Effective degrees of freedom were estimated using the Satterthwaite approximation implemented in *lmerTest*^73^. Finally, to control the family-wise error rate across simple-effect contrasts, *p*-values were adjusted using the Holm-Bonferroni procedure. Statistical analyses of the sensation survey were conducted in Python (version 3.12.3) using the SciPy library^74^. All statistical tests were two-sided, and results were considered significant at *α* = 0.05.

## Acknowledgements

This work was supported by the National Institutes of Health SBIR program (Grants R44NS139730, R44NS127667, 1R44NS127667-01, 1R43MH143230-01, 1R44MH142142-01). Additional support was provided by the Weill Neurohub Next Great Ideas Fellowship and the Weill Neurohub Pillars Program. We thank Owen Doyle for technical development and Tianhe Wang for statistical consultation. Daisy Dai, Saumya Singh, Min Ji (Alicia) Kim, and Havisha Desai assisted with the investigation.

## Author contributions

P.R., C.M., G.A., A.V.P., K.G., L.L., and R.B.I. conceived and designed the study. P.R., C.M., D.S., C.L., L.L., and R.B.I. developed the research tools. P.R. developed the software for experimentation and analysis. I.K., K.T.P., H.A.A., and P.R. collected the data. P.R. performed data curation, formal analysis, E-field simulation, and visualization. C.M., D.S., L.L., and R.B.I. acquired funding. R.B.I. provided laboratory resources. P.R. wrote the original draft. R.B.I. and K.G. edited the manuscript. All authors provided critical review and approved the final paper.

## Data and code availability

Data and analysis code will be made available upon publication.

## Supplementary Materials

**Figure S1.**
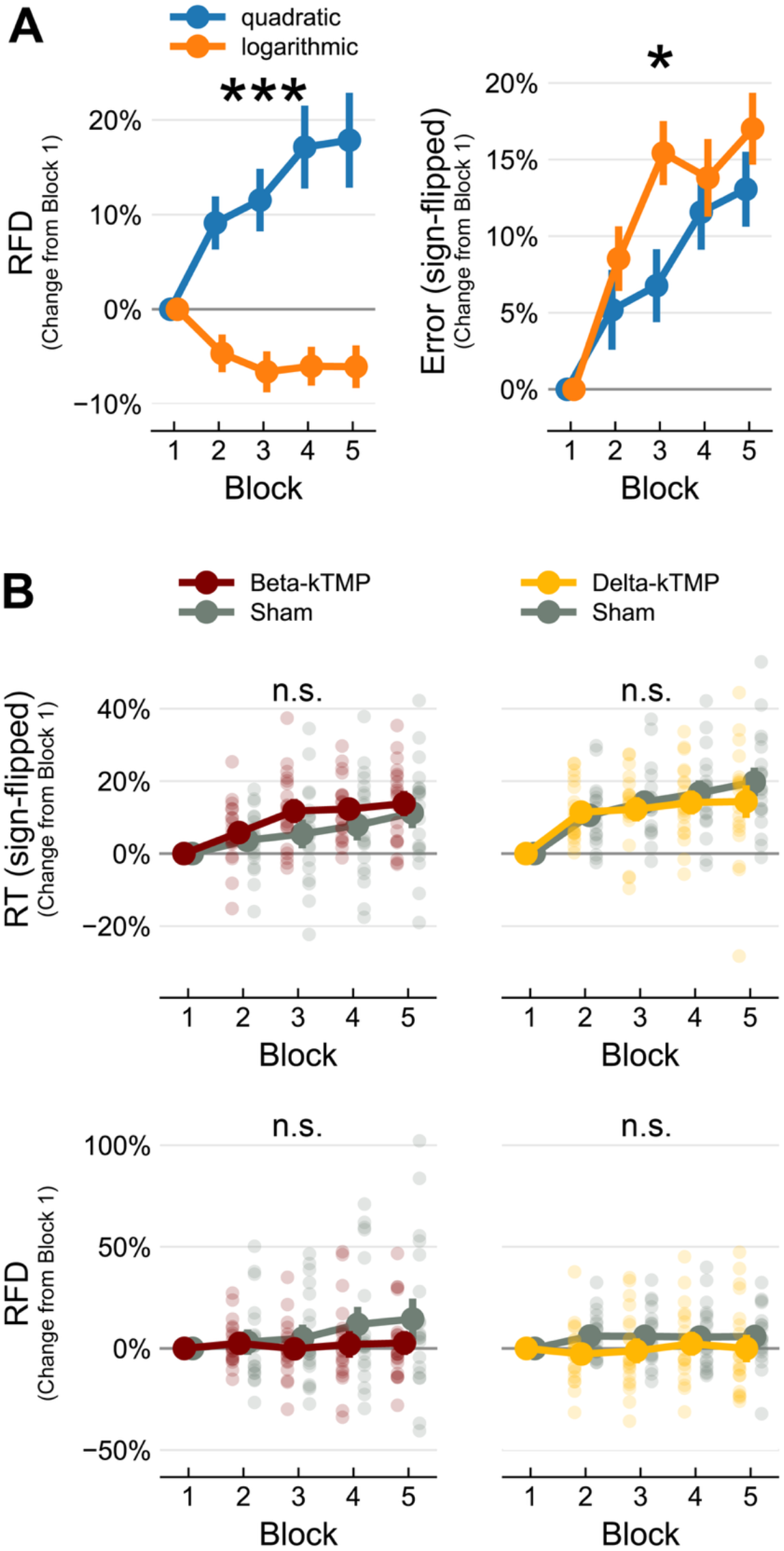
Changes in performance relative to the first block, indexed by the rate of force development (RFD), error, and reaction time (RT). **(A)** Changes in RFD (left) and error (right) differ between force-cursor mappings. **(B)** RT and RFD improve over time, with similar gains during active vs. sham stimulation in either frequency group. All error bars are standard errors.

